# Charged Pore-lining Residues are Required for Normal Channel Kinetics in the Eukaryotic Mechanosensitive Ion Channel MSL1

**DOI:** 10.1101/2020.07.27.222992

**Authors:** Angela M. Schlegel, Elizabeth S. Haswell

## Abstract

Mechanosensitive (MS) ion channels are widespread mechanisms for cellular mechanosensation that can be directly activated by membrane tension. The well-studied MscS family of MS ion channels is found in bacteria, archaea, and plants. MscS-Like (MSL)1 is localized to the inner mitochondrial membrane of *Arabidopsis thaliana*, where it is required for normal mitochondrial responses to oxidative stress. Like *Escherichia coli* MscS, MSL1 has a pore-lining helix that is kinked. However, in MSL1 this kink is comprised of two charged pore-lining residues, R326 and D327. Using single channel patch-clamp electrophysiology in *E. coli*, we show that altering the size and charge of R326 and D327 leads to dramatic changes in open state dwell time. Modest changes in gating pressure and open state stability were also observed while no effects on channel rectification or conductance were detected. MSL1 channel variants had differing physiological function in *E. coli* hypoosmotic shock assays, without clear correlation between function and particular channel characteristics. Taken together, these results demonstrate that altering pore-lining residue charge and size disrupts normal channel state stability and gating transitions, and led us to propose the “sweet spot” model. In this model, the transition to the closed state is facilitated by attraction between R326 and D327 and repulsion between R326 residues of neighboring monomers. In the open state, expansion of the channel reduces inter-monomeric repulsion, rendering open state stability influenced mainly by attractive forces. This work provides insight into how unique charge-charge interactions can be combined with an otherwise conserved structural feature to help modulate MS channel function.

## Introduction

Living organisms constantly experience physical force from both internal and external sources and possess a variety of mechanisms for detecting and responding to key mechanical stimuli (Fruleux et al., 2019; Persat et al., 2015; Yang et al., 2015). Among these mechanisms are mechanosensitive (MS) ion channels, which are found in all kingdoms of life (Hamilton, Schlegel, et al., 2015; Kloda & Martinac, 2001; Kung et al., 2010; Ranade et al., 2015). Most MS channels are opened (gated) primarily by increases in lateral membrane tension (Cox et al., 2019).

While MS ion channels are united by their primary gating stimulus rather than a common mechanosensory sequence or structure, individual MS channel families have been identified by the presence of conserved domains. One such family is the MscS family, which is defined by similarity to the *E. coli* <u>M</u>echano<u>s</u>ensitive ion channel of <u>S</u>mall conductance (*Ec*MscS) (Haswell, 2007; Malcolm & Maurer, 2012; Pivetti et al., 2003). *Ec*MscS, along with the <u>M</u>echano<u>s</u>ensitive ion <u>c</u>hannel of <u>L</u>arge conductance (MscL), allow *E. coli* cells to survive hypoosmotic shock. Sudden transfer into a hypotonic solution leads to water entry into the cell, subsequent swelling, and presumably an increase in lateral membrane tension. Increased membrane tension in turn opens MscS and MscL, allowing for rapid osmoregulation and preventing cell damage (Bialecka-Fornal et al., 2015; Boer et al., 2011; Buda et al., 2016; Levina, 1999; Rojas et al., 2014).

Multiple structures of *Ec*MscS describe a homoheptameric channel with a transmembrane (TM) domain, comprised of three TM helices per monomer, atop a large cytoplasmic “cage” (Bass et al., 2002; Lai et al., 2013; Pliotas et al., 2015; Rasmussen et al., 2019; Reddy et al., 2019; Steinbacher et al., 2007; Wang et al., 2008). A key feature of the *Ec*MscS structure is the pore-lining TM helix, TM3, which, in the nonconducting state, kinks mid-way through at G113, such that its C-terminal portion points outward from the pore and lies parallel to the lipid bilayer (Bass et al., 2002; Lai et al., 2013; Rasmussen et al., 2019; Reddy et al., 2019). During gating, TM3 is proposed to pivot outward around and partially straighten this kink, thus removing pore occlusions and allowing for ion flow (Lai et al., 2013; Pliotas et al., 2015; Vásquez et al., 2008; Wang et al., 2008). Mutations to either G113 or neighboring Q112 alter channel characteristics such as desensitization/inactivation and entry into subconducting states (Akitake et al., 2007; Edwards et al., 2008), highlighting the importance of this structural feature in shaping channel behavior.

Based on homology to the pore-lining domain and top portion of the cytoplasmic domain of *Ec*MscS, MscS family members have been found throughout the bacterial and archaeal kingdoms, in all currently available plant genomes, and in some protist genomes (Basu & Haswell, 2017). The genome of the model flowering plant *Arabidopsis thaliana* encodes ten homologs of *Ec*MscS, termed <u>M</u>sc<u>S</u>-<u>L</u>ike (MSL) channels (Haswell, 2007). MSLs localize to various compartments, including the plasma membrane (Hamilton, Jensen, et al., 2015; Haswell et al., 2008), chloroplast membrane (Haswell & Meyerowitz, 2006), and inner mitochondrial membrane (Lee et al., 2016). Mechanosensitive channel activity has been demonstrated in heterologous systems for MSL1, MSL8, and MSL10 (Hamilton & Haswell, 2017; Lee et al., 2016; Maksaev & Haswell, 2012) and in native membranes for MSL8 and MSL10 (Hamilton, Jensen, et al., 2015; Haswell et al., 2008). MSL2/3 and MSL8 are involved in osmoregulation of chloroplasts and pollen, respectively (Veley et al., 2013; Hamilton, Jensen, et al., 2015; Hamilton & Haswell, 2017), much like *Ec*MscS in *E. coli* cells. However, MSL10 has a cell-death signaling activity that is separable from its MS channel activity (Maksaev et al., 2018; Veley et al., 2014), revealing MSL function beyond maintaining osmotic homeostasis.

MSL1 is localized to the inner membrane of mitochondria and appears to be involved in regulating the redox status of mitochondria during stress (Lee et al., 2016). Of all the Arabidopsis MSLs, it most closely resembles *Ec*MscS in overall structure, channel behavior, and sequence. Structural and biochemical analyses of MSL1 revealed a homoheptameric channel consisting of a TM domain, comprised of 5 TM helices per monomer, atop a large cage region likely to be located in the mitochondrial matrix (Deng et al., 2020; Lee et al., 2016; Li et al., 2020). MSL1 and *Ec*MscS are both slightly anion preferring and have average conductances of ~1.2 nS at negative membrane potentials (Edwards et al., 2008; Lee et al., 2016; Sukharev, 2002). However, compared to *Ec*MscS, MSL1 shows both stronger rectification (a directional preference for ion flow) and hysteresis (a difference in open and closing tensions), with a preference for transporting anions out of the cell, and with channel closure often occurring at lower membrane tension than channel opening (Anishkin et al., 2010; Belyy et al., 2010; Sukharev et al., 2007). A sequence alignment (Figure 1A) revealed strong conservation between the pore-lining helices of MSL1 and *Ec*MscS with a singular exception: two neighboring residues are charged in MSL1 (R326 and D327) and polar in *Ec*MscS (Q112 and G113) (red box, Figure 1A).

**Figure 1.**
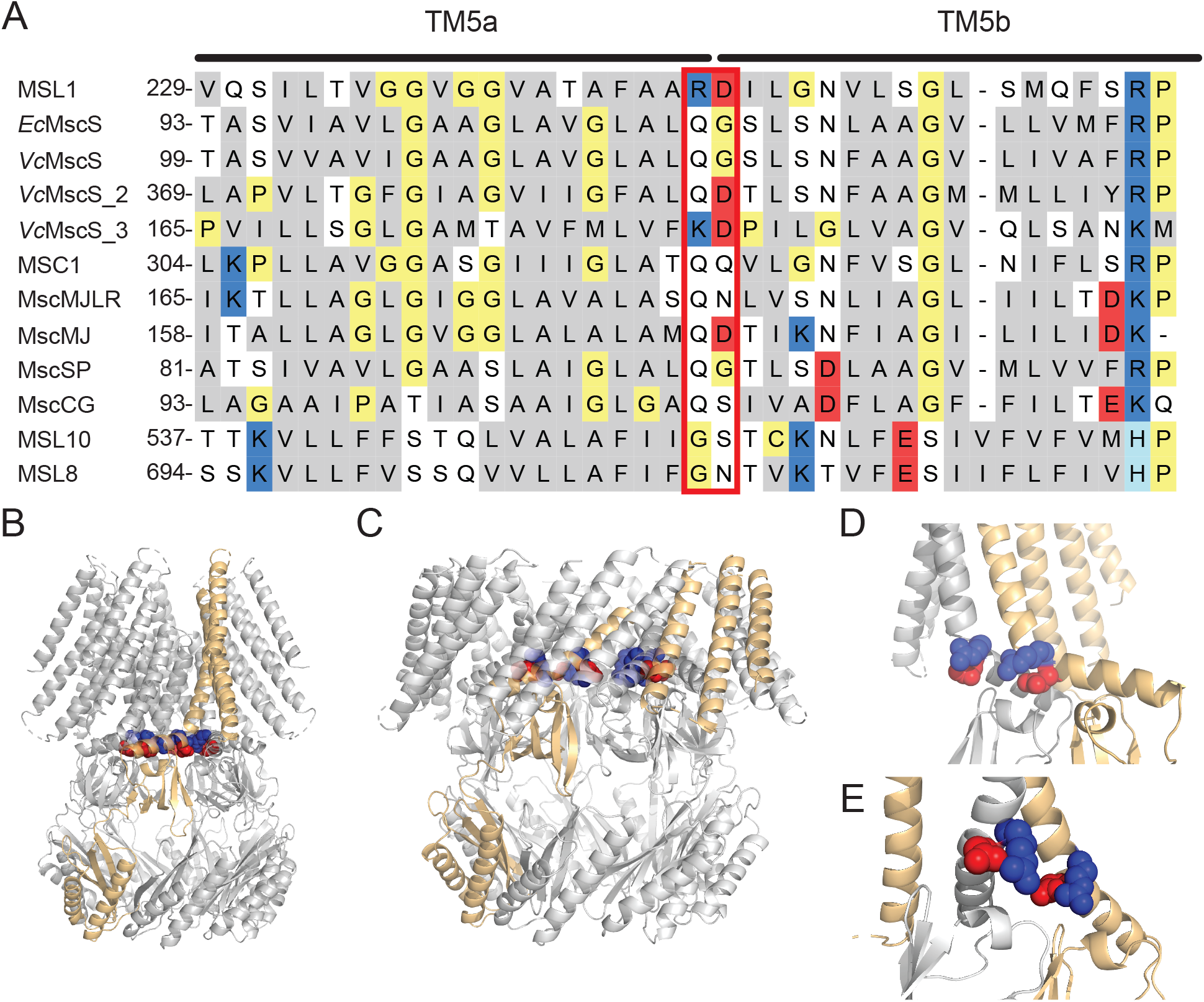
R326 and D327 are charged residues in the kinked pore-lining TM5 helix of the MS ion channel MSL1. (A) Alignment of pore-lining helices from MscS family members for which rectification information is available. Non-polar residues are gray, polar residues white, positively charged residues blue, negatively charged residues red, and other residues pale yellow. R326 and D327 of MSL1 and the corresponding residues in other MscS family members are highlighted by a red box. (B-E) Images of cryoEM structures of MSL1 (PDB file 6VXM (Deng et al., 2020)) and MSL1^A320V^ (PDF file 6VXN (Deng et al., 2020)) in closed and open states, respectively. One monomer is light orange and residues R326 (blue) and D327 (red) are indicated. (B, C) Side view of the placement of R326 and D327 in the TM5 kink of MSL1 (B) and MSL1^A320V^ (C) multimers, respectively. (D, E) Close-up view of the R326 and D327 residues in two adjacent monomers, one grey and one light orange, as viewed from inside the MSL1 (D) and MSL1^A320V^ (E) pores.

Rectification of MSL1 is also strong compared to other MscS family members for which this feature has been characterized (Lee et al., 2016) and most closely resembles that of MscS-like activity detected in *V. cholerae* cells (Rowe et al., 2013). One of three MscS-like genes from *V. cholerae* also encodes a positively charged and a negatively charged residue at the same position as R326 and D327 (Figure 1A). With the exception of MSC1 from *Chlamydomonas reinhardtii* chloroplasts and MscMJ from *Methanocaldococcus jannaschi*, (Kloda & Martinac, 2001; Nakayama et al., 2007), other MscS family members from archaea, bacteria, and plants show only mild rectification (Hamilton, Jensen, et al., 2015; Kloda & Martinac, 2001; Maksaev & Haswell, 2012; Nakayama et al., 2013; Petrov et al., 2013; Edwards et al., 2008). While the correlation between charged residues and rectification in the MscS family is not strict, charged residues have been demonstrated to control rectification in other channels (Li et al., 2008).

Recently reported cryoEM structures of MSL1 in the closed state (Deng et al., 2020; Li et al., 2020) place R326 and D327 at the kink of the pore-lining helix TM5, which is bent such that its C-terminal half runs parallel to the bilayer (Figure 1B), similar to TM3 in the non-conducting state of *Ec*MscS. In the MSL1^A320V^ structure, proposed to represent the open state (Deng et al., 2020), TM5 is almost completely straight and sits diagonally within the bilayer (Figure 1C). These structures support a gating transition in which neighboring R326 and D327 side chains point inward from the TM5 kink in the closed state (Figure 1D), then are pushed towards each other and away from the pore during opening (Figure 1E). TM5 helices from neighboring monomers also move farther apart during channel opening. As with Q112 and G113 of *Ec*MscS (Akitake et al., 2007; Edwards et al., 2008), altering R326 and D327 of MSL1 may affect kink formation and thus channel behavior.

In this study, we investigated the roles of R326 and D327 in MSL1 rectification and other hallmarks of MSL1 channel behavior using single-channel patch-clamp electrophysiology and physiological assays in *E. coli*. Our results provide insight into the roles of individual residues in the MSL1 pore-lining helix and validate recently published MSL1 cryoEM structures (Deng et al., 2020; Li et al., 2020). More broadly, our study contributes to the understanding of how the specific composition of common structural features, like the kinked pore-lining helix found in the MscS family, can influence properties of MS ion channels.

## MATERIALS AND METHODS

### Subcloning and *E. coli* strains

The MSL1 sequence lacking the putative N-terminal mitochondrial transit peptide sequence (residues 1-79; (Lee et al., 2016)), codon-optimized for translation in *E. coli*, was synthesized (ThermoFisher Scientific, USA) and cloned into the pET300 vector to create pET300-MSL1. A C-terminal GFP tag was then added before the stop codon of MSL1 with an EcoRI cut site as the linker sequence between MSL1 and GFP to create pET300-MSL1-GFP. Site directed mutagenesis was then used to create pET300-MSL1^R326Q^-GFP, pET300-MSL1^D327G-GFP^, pET300-MSL1^R326Q D327G-GFP^, pET300-MSL1^D327N^-GFP, and pET300-MSL1^R326Q D327N^-GFP (primer sequences in Table S1). Mutations were verified using restriction enzyme digest and sequencing; the R326Q mutation causes the loss of a PmlI site, the D327G mutation creates an EcoRI site, and the D327N mutation creates a SspI site. To create pET300-MscS-GFP, the MSL1 sequence was replaced with the full-length *Ec*MscS sequence. Lysogenization of *E. coli* strains FRAG-1 (Epstein & Kim, 1971), MJF465 (Levina, 1999), MJF641, and MJF516 (Edwards et al., 2012) was performed using the Novagen λDE3 Lysogenization Kit (Millipore Sigma) following manufacturer’s instructions. Lysogenized strains used in this study are indicated by (DE3).

### Sequence alignment and functional predictions

The MSL1 cryoEM structures (RCSB Protein Data Bank, PDB ID 6VXM (Deng et al., 2020) and 6LYP (Li et al., 2020)) were visualized and images generated using PyMol (Schrödinger, Inc.). MscS family member protein sequences were obtained from publicly available data bases with accession numbers as follows: *Escherichia coli* MscS (*Ec*MscS), UniProt ID P0C0S2; *Arabidopsis thaliana* MSL1 (MSL1), At4g00290; *Arabidopsis thaliana* MSL8 (MSL8), At2g17010; *Arabidopsis thaliana* MSL10 (MSL10), At5g12080; *Corynebacterium glutamicum* MscCG, RefSeq WP_011014245.1; *Chlamydomonas reinhardtii* MSC1, GenBank ID AB288852.1; *Silicibacter pomeroyi* MscSP, UniProt ID Q5LMR6; *Methanococcus jannaschii* MscMJ, UniProt ID Q6M0K6; *M. jannaschii* MscMJLR, UniProt ID Q58543. Structural features of sequences were either assigned based on previously published structural data or, when none was available, predicted using the TMHMM server, v 2.0 (DTU HealthTech). Sequences of 70 amino acids containing predicted or known pore-lining sequences were then aligned in Unipro UGENE software using the built-in MUSCLE algorithm.

### MSL1 variant expression and localization in *E. coli*

Approximately 10 colonies of MJF465(DE3) cells expressing GFP-tagged MSL1 variants were placed into a 14 mL culture tube with 3 mL LB + 1 mM carbenicillin and shaken at 37°C, 250 rpm to an OD_600_ of ~0.5. 2 mL of this culture was added to 100 mL LB + 1 mM carbenicillin and shaken at 37°C, 250 rpm until OD_600_ ~0.5. Isopropyl β-D-1-thiogalactopyranoside (IPTG) was then added to a final concentration of 1 mM and cultures shaken at 37°C, 250 rpm for either 30 min (for expression of MscS-GFP and GFP) or 1 hour (for expression of untagged MSL1 and GFP-tagged MSL1 variants). To image GFP signal, cells were placed on a 1% agarose pad, covered with a coverslip, then imaged using an Olympus FV3000 confocal microscope. GFP was excited using a 488 nm laser and GFP emission was collected from 493-533 nm. For images of cells expressing cytoplasmic GFP, laser transmissivity was 5% and PMT voltage was 436 V. For cells expressing either a GFP-tagged MSL1 variant or MscS-GFP, laser transmissivity was set at 6% and PMT voltage was 515 V. Both bright field and GFP fluorescence images were taken for each sample.

### Patch-clamp electrophysiology

Giant *E. coli* spheroplasts were made according to (Schlegel & Haswell, 2020). The MJF641(DE3) strain was used for conductance analysis, MJF516(DE3) cells for tension sensitivity measurements, and either MJF641(DE3) or MJF516(DE3) cells for open state dwell time measurements. Cells were transformed with the appropriate expression constructs and grown overnight on LB plates containing 1 mM carbenicillin at 37°C. Cells were then cultured in LB with 1 mM carbenicillin at 37°C, 250 rpm to an OD_600_ of 0.4-0.5, then diluted 1:10 in 30 mL LB + 60 μg/mL cephalexin (without carbenicillin) and shaken at 42°C, 180 rpm until cells reached ~75-100 μm in length. IPTG was added to each culture to a final concentration of 1 mM and cultures shaken at 42°C, 180 rpm for 1 hour. Cultures were incubated at 4°C overnight, then spun down at 3000 xg. Cell pellets were gently resuspended in 2.5 mL 0.8 M sucrose and the following spheroplast reaction components added in order to the resuspension, with gentle swirling after each addition: 150 μL 1 M Tris-HCl (pH 7.2), 120 μL 5 mg/mL lysozyme, 50 μL 5 mg/mL DNase I, 150 μL 0.125 M EDTA. The reaction was incubated at room temperature for 5-7 min, then stopped by adding 1 mL stop solution (0.68 M sucrose, 19 mM MgCl_2_, 9.5 mM Tris-HCl pH 7.2, 0.22 μm filter-sterilized) and swirling to mix. 3.5 mL dilution solution (0.78 M sucrose, 1 mM MgCl_2_, 1 mM Tris-HCl pH 7.2, 0.22 μm filter-sterilized) was added, and 275 μL aliquots stored at −80°C.

All data were collected from inside-out configuration patches. The pipette buffer used was 200 mM KCl, 90 mM MgCl_2_, 5 mM CaCl_2_, 5 mM HEPES, pH 7.4. The bath buffer was identical to the pipette buffer with the addition of 400 mM sucrose. Pressure application was controlled using an HSPC-1 pressure clamp system (ALA Scientific Instruments) and data were acquired using an Axopatch 200B amplifier and a Digidata 1440A digitizer (Molecular Devices) at 20 kHz and low-pass filtered at 5 kHz except for open state dwell time measurements, for which data was collected at 10 kHz. Data were analyzed using Clampfit 10.6 (Molecular Devices).

Conductance measurements were performed at membrane potentials ranging from −150 mV to 80 mV using 5 s symmetric pressure ramps. The largest conductance value for each gating event was taken to avoid including potential substate conductance measurements in the average conductance calculations. Conductances were then calculated using Ohm’s law at membrane potentials of −120 mV, −60 mV, and 60 mV.

Tension sensitivity of MSL1 variants was assessed by determining the gating pressure of MSL1 or an MSL1 variant relative to that of endogenously expressed MscL, using 5-10 s symmetric pressure ramps at a membrane potential of −70 mV. The first gating events observed for each channel in a single trace were used and only MSL1 gating events lasting a minimum of 1 s were considered. Data were only analyzed if both MSL1 variant and MscL gating events were observed in the same trace and if no MSL1 variant gating events were observed prior to application of additional negative pressure to the patch.

Open state dwell time measurements were performed using a 2-4 s symmetric pressure ramp followed by monitoring of channel activity until 97.7 s after the start of the pressure ramp. Membrane potential was maintained at −70 mV throughout the course of this protocol. Traces were not analyzed if channel activity was detected prior to application of the pressure ramp and a channel was considered closed if no activity was observed for 5 s. Individual traces were pooled from 10 patches per channel in order to calculate the percentage of gating events falling into one of five open state dwell time bins: 0-19.99 s, 20-39.99 s, 40-59.99 s, 60-79.99 s, 80+ s.

### *E. coli* growth assay

Five freshly transformed MJF465(DE3) colonies were grown at 37°C, 250 rpm in LB with 1 mM carbenicillin to an OD_600_ of ~0.5. Cultures were then diluted to an OD_600_ of 0.05 in either LB only or LB + 1 mM IPTG and three 250 μL aliquots of each dilution transferred to a clear, flat-bottom 96-well plate. This plate was then placed in an Infinite M200 Pro plate reader, then incubated at 37°C with continuous shaking and OD_600_ measurements made every 15 min for a total of 6 h. Growth assays were repeated using cells from three independent transformations.

### *E. coli* hypoosmotic shock survival assay

Assays were conducted as described in (Bartlett et al., 2004) with some modifications. Freshly transformed colonies were grown overnight at 37°C, 250 rpm in low glucose citrate-phosphate media (60 mM Na_2_HPO_4_, 5 mM K_2_HPO_4_, 7 mM citric acid, 7 mM NH_4_SO_4_, 0.4 mM MgSO_4_, 3 μM thiamine, 6 μM iron) with 0.04% glucose and 1 mM carbenicillin. Overnight cultures were diluted 1:5 in citrate-phosphate media with 0.2% glucose and grown to an OD_600_ of ~0.3 at 37°C, 250 rpm. Cultures were then diluted 1:1 in citrate-phosphate media with 0.2% glucose and 1 M NaCl and grown to an OD_600_ of ~0.3, at which point expression was induced for 1 hour by the addition of 1 mM IPTG. Cultures were diluted 1:20 in either ddH_2_O for shocked samples or 0.5 M NaCl citrate-phosphate buffer (60 mM Na_2_HPO_4_, 5 mM K_2_HPO_4_, 7 mM citric acid, 7 mM NH_4_SO_4_) for unshocked controls and shaken at 37°C, 250 rpm for 15 min. Cultures were serially diluted 1:10 six times in either ddH_2_O (shocked samples) or 0.5 M NaCl citrate-phosphate buffer (unshocked controls). A 5 μL aliquot of each dilution was then spotted onto LB + carbenicillin plates and grown overnight at 30°C. The next day, the number of colonies grown from each dilution were counted and survival ratios of shocked/unshocked colonies calculated for each strain/construct combination calculated using values from dilutions producing up to 50 colonies.

## RESULTS

To begin to study the role of R326 and D327 in MSL1 function, an *E. coli* codon-optimized version of MSL1 lacking the predicted mitochondrial target sequence (2-79 aa; (Lee et al., 2016)), was fused to GFP and expressed from the T7-inducible pET300 vector. For all experiments, constructs were transformed into lysogenized *E. coli* containing IPTG-inducible T7 promoters (see Methods). Four different lysogenized *E. coli* strains were used: MJF465(DE3) (*mscS-mscK-mscL-* (Levina, 1999)), MJF516(DE3) (*mscS-mscK-ybiO-yjeP-* (Edwards et al., 2012)), MJF641(DE3) (*mscS-mscK-ybdG-ybiO-yjeP-ynaI-mscL-* (Edwards et al., 2012)), and their parental strain FRAG-1(DE3) (Epstein & Kim, 1971).

### GFP-tagged MSL1 variants localize to the periphery of *E. coli* cells and do not affect cell growth

We assessed the expression and localization of GFP-tagged MSL1 variants in *E. coli* strain MJF465(DE3) cells by imaging induced cells using a confocal microscope (Figure 2A). All versions of GFP-tagged MSL1 produced punctate GFP signal around the cell periphery that was similar to *Ec*MscS-GFP (as previously observed (Romantsov et al., 2010; van den Berg et al., 2016)), and distinct from cytoplasmic free GFP. Growth rates of all strains were indistinguishable with (Figure 2B) or without (Figure 2C) IPTG (Okada et al., 2002).

**Figure 2.**
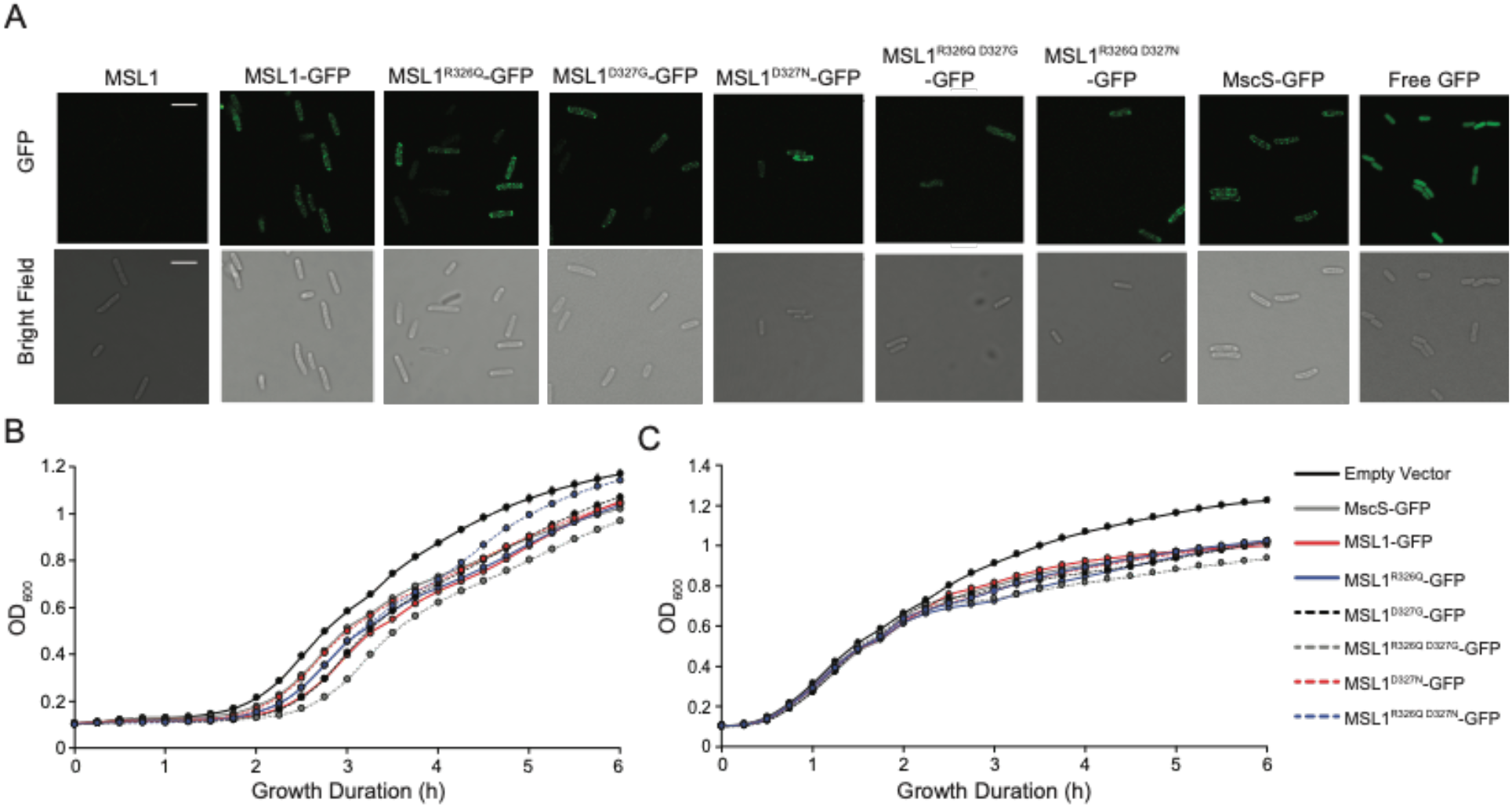
MSL1 variants localize to *E. coli* cell membranes and do not impact *E. coli* cell growth in LB. (A) Confocal micrographs of MJF465(DE3) cells expressing untagged MSL1, MSL1-GFP, a GFP-tagged MSL1 variant, MscS-GFP, or cytoplasmic GFP. Scale bars are 5 μm. (B-C) Growth curves of MJF465(DE3) cells transformed with pET300 vectors encoding the indicated protein or an empty pET21b(+) control. Cells were grown in LB with (B) or without (C) IPTG and OD_600_ values measured every 15 min. Data points are shown ± standard deviation, although error bars may be too small to be visible.

### Mutations to R326 and D327 do not alter channel conductance or rectification

We next sought to characterize the channel behavior of MSL1-GFP variants using single-channel patch-clamp electrophysiology in giant *E. coli* spheroplasts as in (Schlegel and Haswell, 2020). IV curves with membrane potentials ranging from −150 mV to 80 mV for each GFP-tagged MSL1 variant are shown in Figure 3. As demonstrated previously (Lee et al., 2016), MSL1-GFP channel activity was triggered by application of suction to inside-out excised patches and was characterized by a single-channel conductance of ~1.2 nS at negative membrane potentials and markedly reduced conductance at membrane potentials greater than 20 mV. No major differences were observed between the IV curves of MSL1-GFP and any GFP-tagged MSL1 variant. Thus, none of the mutations to R326 nor D327 we tested changed the rectification behavior of MSL1.

**Figure 3.**
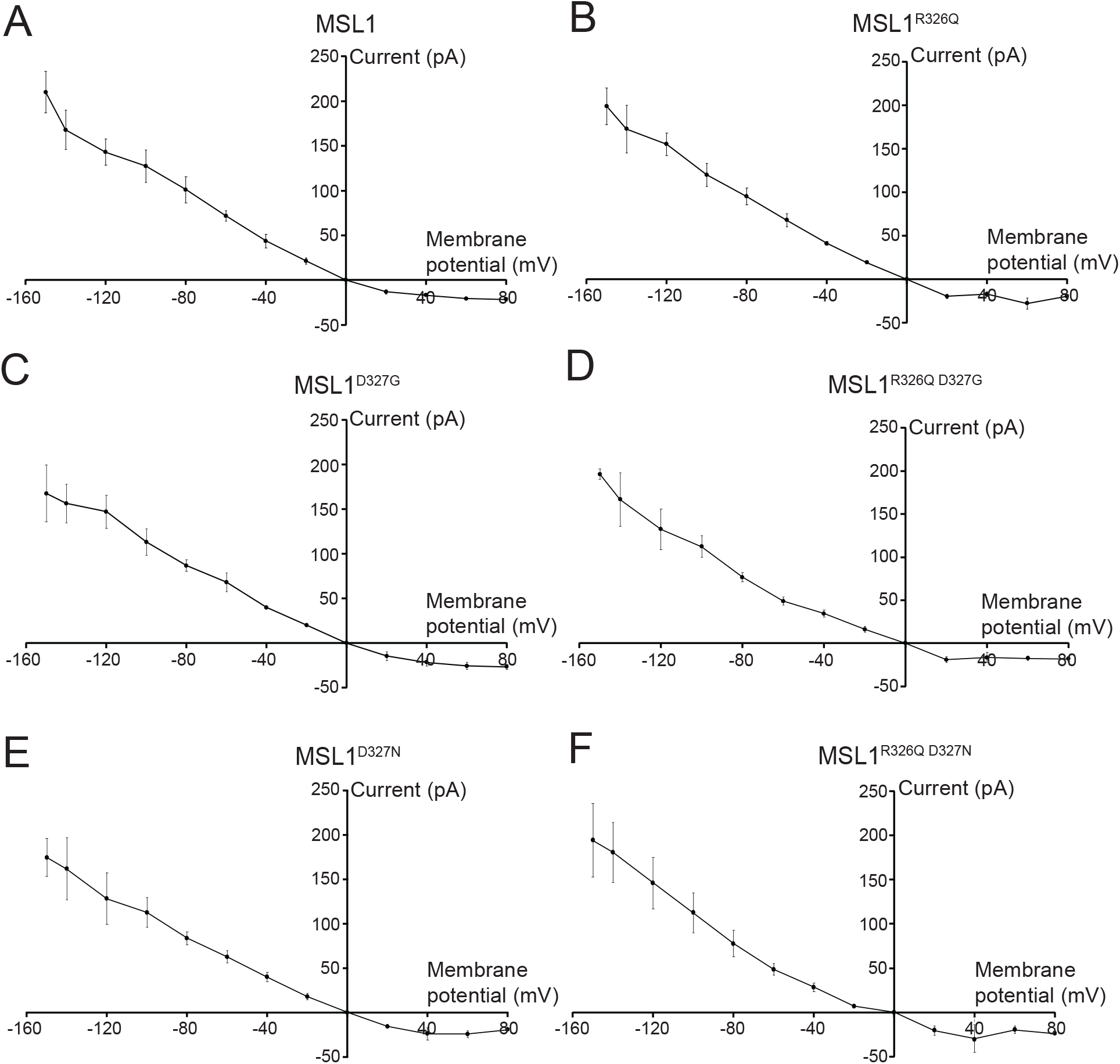
Mutations to R326 and D327 of MSL1 do not affect rectification. IV curves for GFP-tagged MSL1 variants expressed in MJF641(DE3) cells. Each data point represents the average single-channel current for 3 to 17 patches. Error bars indicate standard deviation.

The IV curves shown in Figure 3 were used to calculate conductance at 60 mV, −60 mV, and −120 mV for each GFP-tagged MSL1 variant (Table 1). The single-channel conductances of MSL1^R326Q D327G^-GFP and MSL1^R326Q D327N^-GFP were significantly lower than that of MSL1-GFP at −60 mV (0.82±0.08 nS, 0.81±0.11 nS, and 1.19±0.10 nS, respectively). However, no significant differences in conductance between any variants were detected at 60 mV nor −120 mV. Conductances at - 120 mV are the most physiologically relevant, as plant mitochondria maintain very negative inner membrane potentials (Gerencser et al., 2012; Schwarzländer et al., 2012). In a previous characterization of MSL1^R326Q D327G^, (Li et al., 2020) reported a reduced single channel current but greater total current than the wild type. While these results were interpreted as a higher number of channels open, they could also be due to longer open state dwell times (see below). Taken together, the data shown in Figure 2, Figure 3, and Table 1 indicate that the size and charge at 326 and 327 are not critical for protein stability, localization, or single channel conductance. Unexpectedly, changing R326 and D327 to the analogous resides in *Ec*MscS did not reduce MSL1 rectification (Figure 3).

**Table 1.**
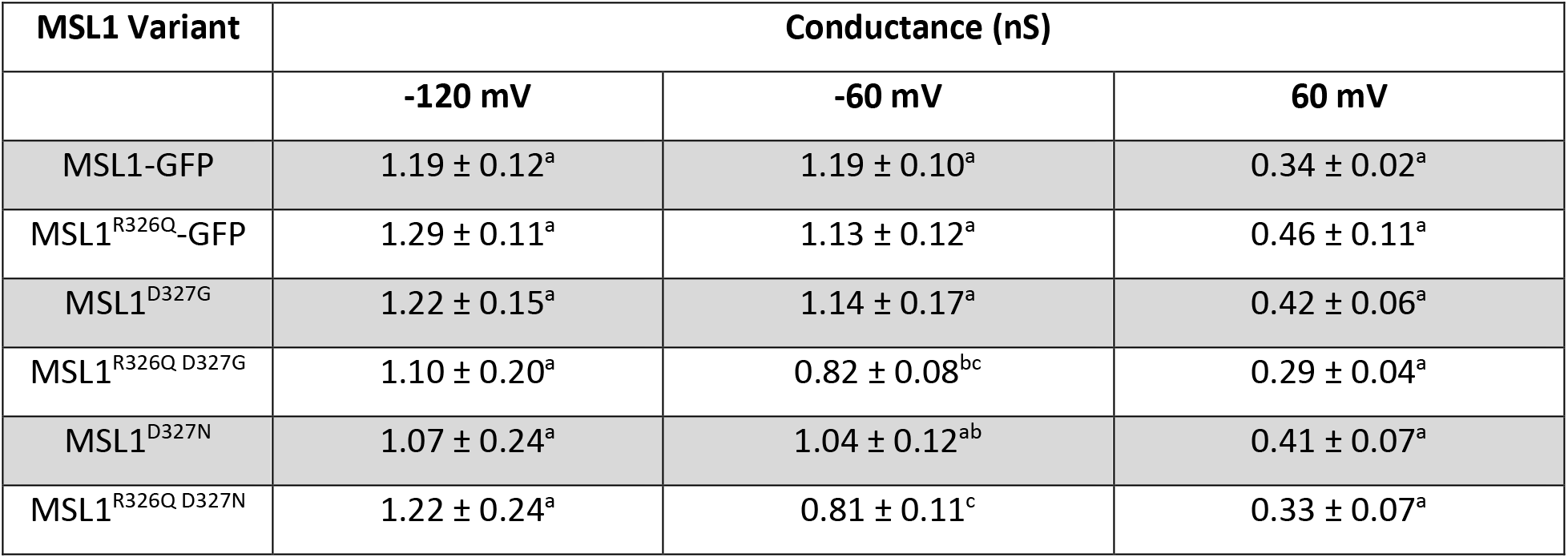
Mutations to R326 and D327 in MSL1 have little effect on channel conductance. Conductance values represent the mean of average patch conductances for 3-7 patches per variant. Differences were statistically evaluated using one-way ANOVA with post-hoc Scheffe’s test; letters indicate statistical differences (p < 0.05).

| MSL1 Variant | Conductance (nS) |  |  |
| --- | --- | --- | --- |
|  | -120 mV | -60 mV | 60 mV |
| MSL1-GFP | 1.19 ± 0.12<sup>a</sup> | 1.19 ± 0.10<sup>a</sup> | 0.34 ± 0.02<sup>a</sup> |
| MSL1<sup>R326Q</sup>-GFP | 1.29 ± 0.11<sup>a</sup> | 1.13 ± 0.12<sup>a</sup> | 0.46 ± 0.11<sup>a</sup> |
| MSL1<sup>D327G</sup> | 1.22 ± 0.15<sup>a</sup> | 1.14 ± 0.17<sup>a</sup> | 0.42 ± 0.06<sup>a</sup> |
| MSL1<sup>R326Q D327G</sup> | 1.10 ± 0.20<sup>a</sup> | 0.82 ± 0.08<sup>bc</sup> | 0.29 ± 0.04<sup>a</sup> |
| MSL1<sup>D327N</sup> | 1.07 ± 0.24<sup>a</sup> | 1.04 ± 0.12<sup>ab</sup> | 0.41 ± 0.07<sup>a</sup> |
| MSL1<sup>R326Q D327N</sup> | 1.22 ± 0.24<sup>a</sup> | 0.81 ± 0.11<sup>c</sup> | 0.33 ± 0.07<sup>a</sup> |

### Mutations to R326 and D327 have modest effects on MSL1 tension sensitivity

Given that R326 and D327 did not affect rectification, we next wished to examine their role in the gating process of MSL1. We started by determining the gating pressure of each MSL1-GFP variant. Gating pressure is a proxy for tension sensitivity; for MS channels in *E. coli* it is often measured relative to endogenously expressed MscL and reported as the pressure threshold ratio (P_x_/P_L_) (Blount et al., 1996). We expressed each GFP-tagged MSL1 variant in *E. coli* strain MJF516(DE3) (Edwards et al., 2012) and generated giant spheroplasts. Using 5-10 s pressure ramps, we measured gating pressures of the first channel openings of each GFP-tagged MSL1 variant and of MscL, and calculated the P_x_/P_L_ values for each variant (Figure 4). MSL1^R326Q D327G^-GFP, MSL1^D327N^-GFP, and MSL1^R326Q D327N^-GFP had significantly higher P_x_/P_L_ than MSL1-GFP (0.65-0.71 compared to 0.49, respectively). In contrast, pressure threshold ratios of MSL1^R326Q^-GFP, MSL1^D327G^-GFP, and MSL1-GFP could not be statistically distinguished, although the average P_x_/P_L_ of individual patches containing MSL1^D327G^-GFP were typically lower than those of MSL1-GFP. These results thus indicate that both size and charge at the MSL1 TM5 kink influence gating pressure, and that the residue at 327 appears to play a dominant role.

**Figure 4.**
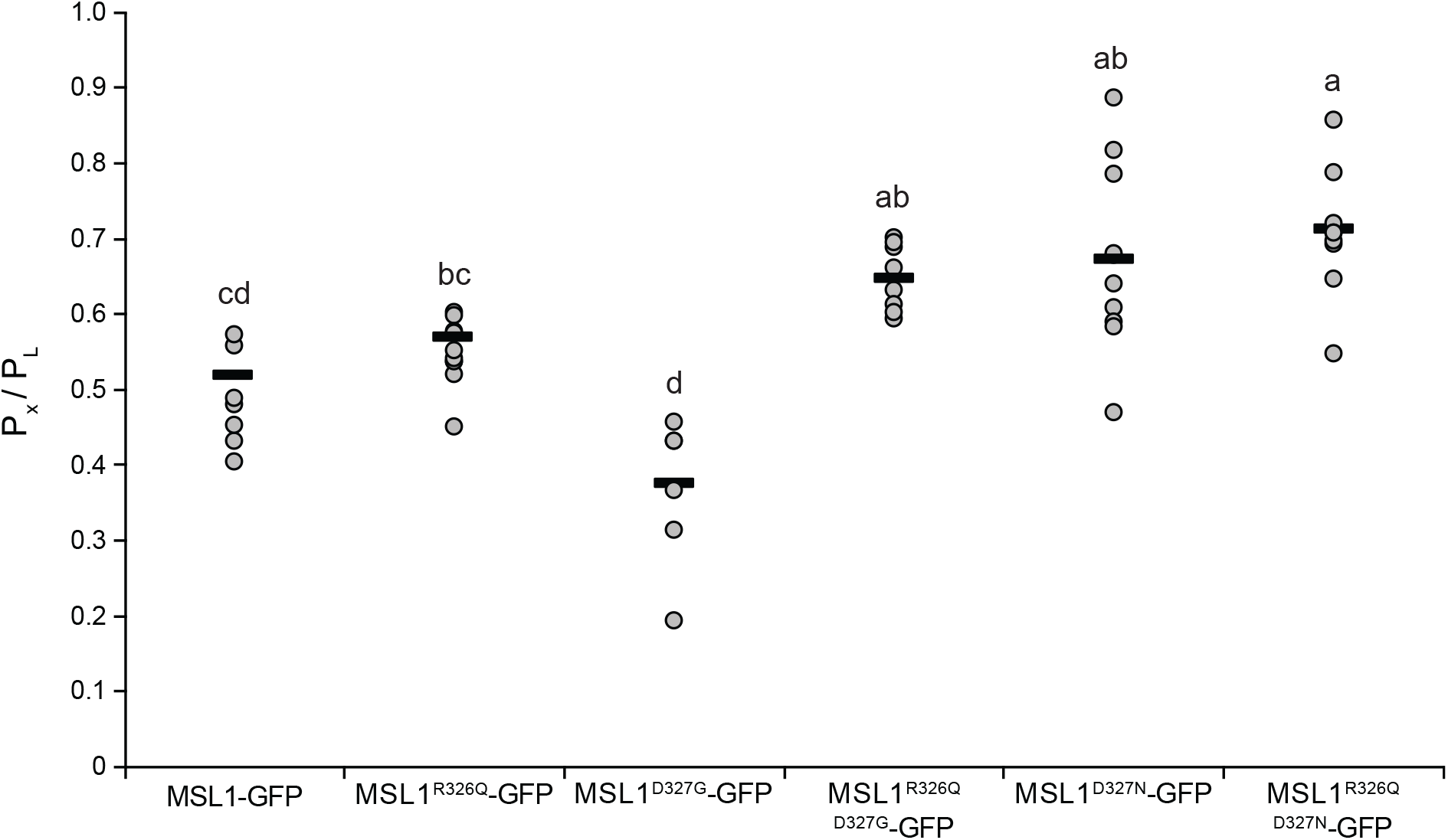
MSL1^R326Q D327G^-GFP, MSL1^D327N^-GFP, and MSL1^R326Q D327N^-GFP have significantly higher gating pressures than MSL1-GFP. Gating pressures of the indicated GFP-tagged MSL1 variants relative to the gating pressures of endogenously expressed MscL. Channels were gated using 5-10 s symmetric pressure ramps at a membrane potential of −70 mV. Each gray circle represents the average of all gating pressure ratios obtained for a single patch, while the black bars represent the mean of patch averages for each sample. N = 6-10 patches per variant. Statistical differences were examined using one-way ANOVA with post-hoc Scheffe’s test; significant differences are indicated by different letters (p < 0.05). Data points greater than two standard deviations beyond the sample average were excluded.

### R326 and D327 exert dramatic and opposing effects on open state dwell time

We also examined the open state dwell times of GFP-tagged MSL1 variants (Figure 5). Using a modified version of a previously published protocol (Akitake et al., 2007), mechanosensitive gating was triggered by applying a brief 2-4 s negative pressure ramp, then the same membrane potential of −70 mV was maintained without any additional suction for a total of 100 s as in (Deng et al., 2020). We then recorded the time from the initial pressure-triggered channel opening to final channel closure, defined as complete cessation of channel activity for 5 s (Figure 5). Most (89%) of MSL1-GFP channel openings lasted less than 20 s, and only 5.5% lasted for more than 80 s. 100% of MSL1^R326Q^-GFP channel openings lasted less than 20 s. In contrast, a large proportion of MSL1^D327G^-GFP and MSL1^D327N^-GFP, channel openings lasted for more than 80 s (62.5% and 72.9%, respectively). Adding the R326Q mutation to these channels reduced the proportion of extremely long open dwell times to 48.4% and 42.1% for MSL1^R326Q D327G^-GFP and MSL1^R326Q D327N^-GFP, respectively (Figure 5). To summarize, we found that reducing the size and positive charge of the amino acid at position 326 decreased open dwell time, reducing the size and negative charge of position 327 amino acid increased open dwell time, and double mutants showed an intermediate open dwell time, suggesting that R326 and D327 in TM5 of MSL1 have opposite effects on closure efficiency.

**Figure 5.**
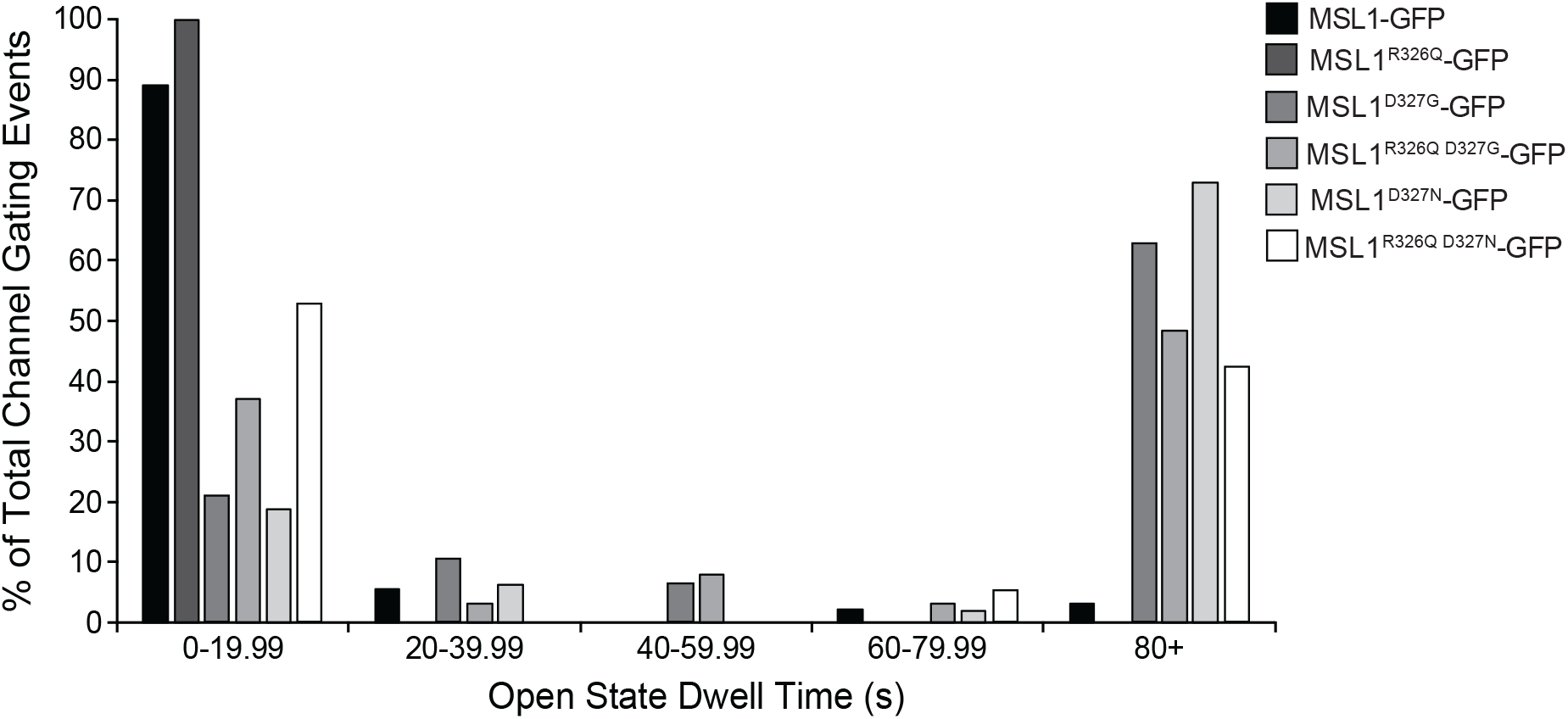
Effect of R326 and D327 mutations on the open state dwell time of MSL1-GFP variants. Membrane potential was maintained at −70 mV and channel gating was triggered by either a 2 s or 4 s symmetric pressure ramp followed by monitoring of channel activity without additional pressure until 97.7 s. Results from 19-97 traces from 9-10 patches per variant are shown.

### Some MSL1 variants have unstable open states

Individual traces (Figure 6) at both −60 mV and −120 mV showed generally stable open states for MSL1-GFP, MSL1^R326Q^-GFP, and MSL1^D327N^-GFP. However, MSL1^R326Q D327G^-GFP, MSL1^R326Q D327N^-GFP, and MSL1^D327G^-GFP were flickery (Figure 6). Flickery channel behavior is produced by rapid transitions between nonconducting, conducting, and subconducting states, and is thought to be indicative of an unstable open state (Malcolm & Blount, 2015; Rasmussen et al., 2007). Thus, both the size and charge of residues at 326 and 327 are important to the stability of the MSL1 open state.

**Figure 6.**
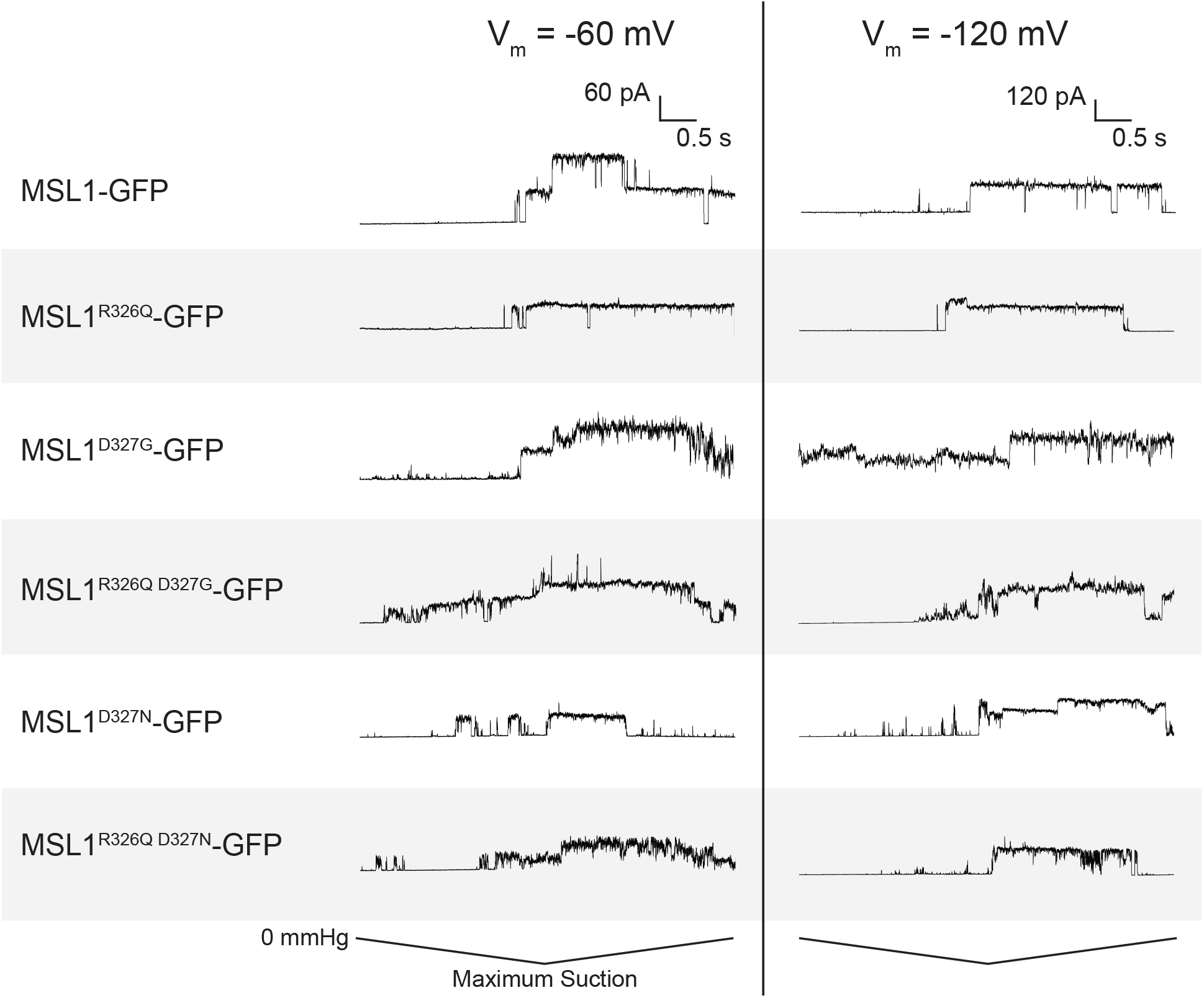
R326 and D327 influence open state stability of MSL1. Representative traces from inside-out excised patches showing pressure-activated gating events of MJF641(DE3) cells expressing the indicated constructs at two membrane potentials. Traces show current measurements taken during a 5 s symmetric negative pressure ramp, with the maximum amount of negative pressure (and therefore rate of pressure application) varying between traces.

### R326 and D327 mutations alter the physiological function of MSL1 in *E. coli*

Like *Ec*MscS, MSL1 provides protection from hypo-osmotic shock to *E. coli* (Lee et al., 2016). To determine the effects of R326 and D327 mutations on this osmoregulatory function, we examined the ability of *E. coli* MJF465(DE3) cells expressing GFP-tagged MSL1 variants to survive hypoosmotic shock. MJF465(DE3) cells lack MscS, MscL, and MscK and therefore cannot survive severe hypoosmotic shock without expressing a functional MS ion channel (Levina, 1999). In this assay, cells are grown in high salt citrate-phosphate media, channel expression is induced, then cells are either hypoosmotically shocked in water or transferred to the same high salt media. FRAG-1(DE3) cells, which contain all endogenous MS channels, survive, while MJF465(DE3) cells do not. MSL1-GFP, MSL1^R326Q^-GFP, and MSL1^R326Q D327G^-GFP all conferred hypoosmotic shock survival rates comparable to that of FRAG-1 cells, suggesting they all contribute to osmoregulation during hypoosmotic shock (Figure 7A, B). Survival rates conferred by MSL1^D327G^-GFP expression were unusually variable and often higher for shocked cells than nonshocked cells (average survival rate of 160%, Figure 7A). Cells expressing MSL1^D327N-GFP^ or MSL1^R326Q D327N-GFP^ grew too slowly in citrate-phosphate media to be analyzed in this assay.

**Figure 7.**
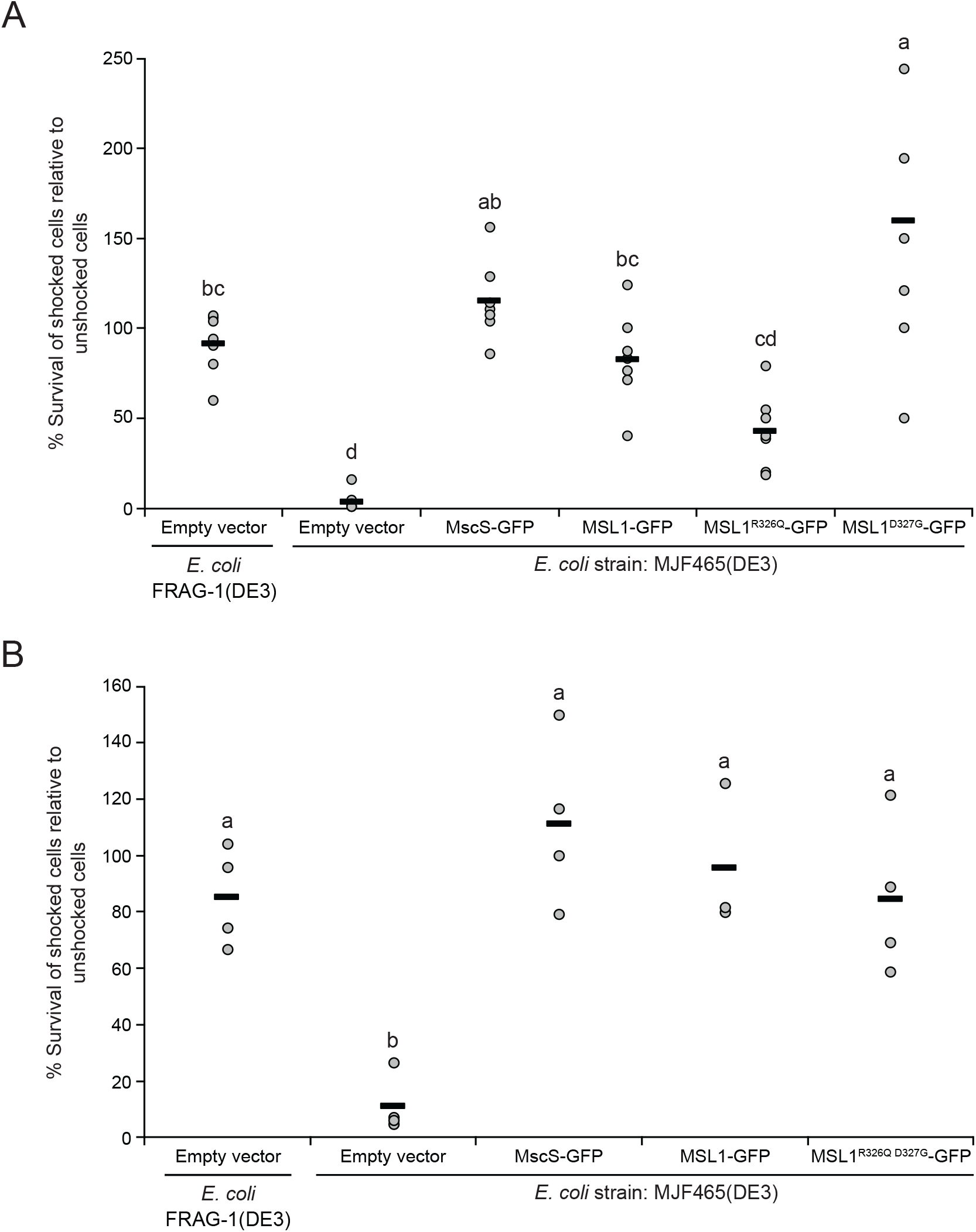
Some MSL1 variants protect *E. coli* strain MJF465(DE3) from hypoosmotic shock. Hypoosmotic shock survival rates of cells from the indicated strains relative to unshocked controls. Each circle represents the relative survival rate for an experiment and black bars indicate the average survival rate for all experiments. For each panel, statistical differences were evaluated using one-way ANOVA followed by a post-hoc Scheffe’s test; different letters indicate significant differences (p < 0.05). One data point greater than two standard deviations beyond the sample average was excluded.

MSL1-GFP variants thus had a variety of effects on *E. coli* physiology that may be attributed to a combination of gating pressure (Figure 4), open state dwell time (Figure 5), and open state stability (Figure 6). The reduced open dwell time of MSL1^R326Q^-GFP and extended open dwell time and increased gating pressure of MSL1^R326Q D327G^-GFP did not seem to affect their function in *E. coli* cells during hypoosmotic shock. In contrast, MSL1^D327G^-GFP provided large variations in protection between experiments, perhaps due to the combination of a lower gating threshold and extended open dwell times. It is unclear from our electrophysiological analysis why MSL1^D327N-GFP^ and MSL1^R326Q D327N^-GFP impaired cell growth, as they had higher gating pressures than MSL1-GFP and therefore do not fit classic gain-of-function characteristics (Blount et al., 1997).

## DISCUSSION

The *Arabidopsis* mitochondrial MS channel MSL1 contains a notable feature midway through its pore-lining TM5 helix: a kink formed by charged residues R326 and D327. In *Ec*MscS, the pore-lining kink is proposed to play important roles in transitions between channel states (Akitake et al., 2007; Edwards et al., 2008; Lai et al., 2013; Pliotas et al., 2015; Vásquez et al., 2008; Wang et al., 2008), but the residues that comprise it are nonpolar. To determine the role played by R326 and D327 in both distinct and shared characteristics of MSL1 and *Ec*MscS, we created MSL1 variants in which the charges and size of R326 and D327 were altered, then evaluated their channel behavior and physiological function in *E. coli* Mutations to R326 and D327 affected tension sensitivity, open state dwell time, and open state stability, indicating a role in modulating MSL1 channel state stabilities and transitions, but did not affect stability, localization, conductance, nor rectification.

Based on open and closed state cryoEM structures, we have proposed that MSL1 opening is driven by membrane flattening and area expansion (Deng et al., 2020). These forces drive the outward rotation and tilting of TM5 and the straightening of the kink that joins TM5a and TM5b during the MSL1 gating transition. The data presented here, summarized in Table 2, suggest that the charge and size of R326 and D327 side chains are important for the stability of the open state and for gating and closing transitions. Combining these results with cryoEM structures (Deng et al., 2020; Li et al., 2020), we infer that in the closed state, charge-charge repulsion between R326 side chains on different monomers is finely balanced by charge-charge attractions between R326 and D327 within each monomer (Figure 1B, D). In the open state, intra-monomeric attractive forces between R326 and D327 dominate and inter-monomeric repulsions lose strength, due to the increased distance between helices from different monomers and the shortened distance between R327 and D327 (Figure 1C, E). Below, we describe how our results can be explained by this “sweet spot” model.

**Table 2.** Summary of GFP-tagged MSL1 variant properties. Conductance and gating pressure are presented relative to MSL1-GFP measurements. ns indicates differences from WT are not statistically significant.

| MSL1 Variant | Conductance | Gating Pressure | Open State Stability | Open State Dwell Time |
| --- | --- | --- | --- | --- |
| WT MSL1 | - | - | Stable | - |
| MSL1<sup>R326Q</sup> | WT | 1.12 WT<sup>ns</sup> | Stable | Short |
| MSL1<sup>D327G</sup> | WT | 0.75 WT<sup>ns</sup> | Flickery | Very Long |
| MSL1<sup>R326Q D327G</sup> | Low at -60 mV | 1.32 WT | Slight Flicker | Long |
| MSL1<sup>D327N</sup> | WT | 1.39 WT | Stable | Very Long |
| MSL1<sup>R326Q D327N</sup> | Low at -60 mV | 1.45 WT | Slight Flicker | Long |

The most dramatic effect of the lesions we created was on open dwell time, where MSL1^D327G^-GFP, MSL1^R326Q D327G^-GFP, MSL1^D327N^-GFP, and MSL1^R326Q D327N^-GFP variants stayed open for much longer times than MSL1-GFP (Figure 5). We interpret this to reflect the difficulty of the closing transition. All mutations to D327 had a longer open dwell time, suggesting that the charge-charge attraction between D327 and R326 facilitates closure. In contrast, MSL1R326Q-GFP exhibited decreased open dwell time (Figure 5). According to our sweet spot model, the R326Q mutation on its own also would suffer from a loss of charge-charge attraction, but this effect is overshadowed by the loss of repulsion between R326 on different monomers in the closed state. Combining mutations in both residues leads to a channel where both attractive and repulsive forces are lost, and the dwell time is intermediate between the two single mutants. A seemingly counterintuitive observation is that three channels (MSL1^D327G^-GFP, MSL1^R326Q D327G^-GFP, and MSL1^R326Q D327N^-GFP) have both long open dwell times and are flickery. Perhaps these channels have both an unstable open state (hence the flickering) and an increased barrier to closing. Once they are stably closed, however, they stay closed until additional tension is applied.

Modest but statistically significant increases in gating pressure were observed with MSL1^R326Q D327G^-GFP, MSL1^D327N^-GFP, and MSL1^R326Q^ D327N-GFP (Figure 4). These results cannot be easily explained by the sweet spot model described above, but are reminiscent of the attractive charge-charge interactions between the transmembrane and cytoplasmic domains of *Ec*MscS (Machiyama et al., 2009; Nomura et al., 2008). We also observed a mild decrease in the gating pressure of MSL1^D327G^-GFP (Figure 4). This may arise from destabilization of the closed state due to the loss of attractive charge-charge interactions and dominance of repulsive forces. The addition of the R326Q mutation in the MSL1^R326Q D327G^-GFP may ameliorate this closed state repulsion, reversing the effects of the D327G mutation (Figure 4). However, due to the subtlety of all gating pressure changes we observed, other factors may also play a role that are beyond the scope of our model.

The results presented here establish the importance of two rings of oppositely charged neighboring residues in the channel pore in modulating channel kinetics and open state stability for the mitochondrial MS ion channel MSL1. Our data support a sweet spot model wherein attraction between oppositely charged residues on the same monomer and repulsion from identical residues on different monomers work together to facilitate opening and closing transitions as well as the stability of the closed and open states. Given their position at the pore-lining helix kink, a structural feature with demonstrated importance in *Ec*MscS gating (Akitake et al., 2007; Edwards et al., 2008), this work provides a glimpse into how the same structural features can be composed of entirely distinct residues amongst members of the same MS channel family, creating different mechanisms of control. These results provide a starting point for future investigations into the fine-tuning of MSL1 gating transition, as well as insight into the dynamic network of side chain interactions contributing to MS channel behavior.

## ACKNOWLEDGMENTS

We are grateful to Haswell Lab members past and present for valuable comments, support, and training. This work was supported by and HHMI-Simons Faculty Scholar Grant #55108530 to E.S.H.A.M.S. was supported by funding from the Spencer T. and Ann W. Olin Fellowship for Women in Graduate Study and the William H. Danforth Plant Sciences Fellowship

