## Supplemental Table for "Charged Pore-lining Residues are Required for Normal Channel Kinetics in the Eukaryotic Mechanosensitive Ion Channel MSL1"

| Primer Name | Sequence | Template |
|---|---|---|
| R326Q_F | ATGGCCGGAGTTAGGTTATCGC | MSL1 |
| R326Q_R | CTTCTCCCTATCAAGCCCACC | MSL1 |
| D327G_F | CAAGAAGACGACGATGGCTGT | MSL1 |
| D327G_R | ATATTGACCATCATACTCATTGC | MSL1 |
| R326Q_D327G_F | GCATTTGCAGCACAGGGAATTCTGGGTAATG | $MSL1^{R326Q}$ |
| R326Q_D327G_R | CATTACCCAGAATTCCCTGTGCTGCAAATGC | $MSL1^{R326Q}$ |
| D327N_F | CCGCATTTGCAGCACGTAATATTCTGGGTAATG | MSL1 |
| D327N_R | CATTACCCAGAATATTACGTGCTGCAAATGCGG | MSL1 |
| R326Q_D327N_F | CCGCATTTGCAGCACAGAATATTCTGGGTAATG | $MSL1^{R326Q}$ |
| R326Q_D327N_R | CATTACCCAGAATATTCTGTGCTGCAAATGCGG | $MSL1^{R326Q}$ |

**Table S1. Site-directed mutagenesis primers.** Primers used to generate MSL1 variants, their corresponding sequences, and the template used.
